# BAG3 modulates clathrin-mediated endocytosis and tau uptake in astrocytes

**DOI:** 10.64898/2025.12.17.694879

**Authors:** Joel Rodwell-Bullock, Ella Blau, Archan Ganguly, Carol Deaton, Gail V.W. Johnson

**Affiliations:** Departments of Anesthesiology and Perioperative Medicine, Rochester NY, 14620 USA; Departments of Neuroscience University of Rochester, Rochester NY, 14620 USA

## Abstract

In a healthy brain, astrocytes maintain neuronal homeostasis by removing proteins like tau from the extracellular space, preventing their uptake by neurons. In Alzheimer’s disease (AD), this function is impaired, contributing to tau accumulation. Many AD risk genes are linked to endocytosis pathways, suggesting their role in AD pathogenesis. Astrocytes can internalize, degrade, and release tau, but the mechanisms remain unclear. BAG3, a multifunctional protein, regulates vacuolar processes and interacts with clathrin-mediated endocytosis (CME) components. However, its role in astrocytic CME and tau processing is not fully understood. We show BAG3 depletion in astrocytes reduces clathrin-AP-2 interaction, inhibits CME-dependent epidermal growth factor receptor internalization, and decreases tau uptake. Live-cell imaging reveals BAG3 depletion impairs CME dynamics, increasing clathrin particle lifetimes. BAG3 depletion also alters endolysosomal compartments, increasing Lamp1+ puncta and tau co-localization. These findings highlight BAG3’s role in CME, tau trafficking, and vacuolar processes, suggesting its dysfunction may contribute to AD pathogenesis.

## INTRODUCTION

Astrocytes play an indispensable role in supporting neuron health through a variety of mechanisms, including maintenance of the extracellular environment’s composition and organization (Liu et al., 2025). Recent studies have provided evidence that astrocytes can protect against the progression of tau pathology in Alzheimer’s disease (AD) and other tauopathies by taking up extracellular tau before it can be internalized by adjacent neurons, thus preventing spread of neuronal tau burden (Perea et al., 2019; Wang and Ye, 2021). However, defects in the endocytic processes can limit the ability of astrocytes to take up tau and properly clear it (Eisenbaum et al., 2024; Narayan et al., 2020), and contribute to pathogenesis by facilitating the spreading of pathological tau species (Mothes et al., 2023). Indeed, there is significant biological and pathological evidence that dysfunction of endolysosome pathways occurs at the earliest stages of the development of AD (Cataldo et al., 2000; Colacurcio et al., 2018; Ginsberg et al., 2010). Additionally, GWAS have identified many AD-associated risk genes that are key players in the endolysosome system (Szabo et al., 2022). Of note, many of these AD-associated risk genes in the endolysosome pathway are involved in clathrin-mediated endocytosis (CME) (Ando et al., 2020). Therefore, understanding the regulation of CME in astrocytes, and the factors that may contribute to CME dysfunction, is of fundamental importance.

A key modulator of vacuolar dependent processes and tau clearance is the multidomain co-chaperone Bcl-2-Associated Athanogene 3 (BAG3) (Lei et al., 2015; Lin et al., 2023; Lin et al., 2024; Lin et al., 2022b; Sheehan et al., 2023). Using an unbiased proteomic analysis, we previously demonstrated that BAG3 interacts with numerous proteins that regulate endolysosome pathways. We have also specifically demonstrated that, in neurons, BAG3’s interaction with TBC1D10B, a GTPase activating protein for Rab35, facilitates ESCRT-dependent processes and tau clearance (Lin et al., 2022b). Another key group of BAG3 interactors are those involved in CME, including clathrin heavy chain, dynamin, and Ap2a1 and Ap2a2 (members of the AP-2 complex) (Lin et al., 2022b; Liu et al., 2021; Szabo et al., 2022). These data suggest that the effect on tau may occur through multiple separate, but interrelated, pathways.

Given the importance of astrocytic endocytosis in general and, more specifically, astrocytic CME, in maintaining a neurosupportive environment within the brain (Jaye et al., 2024; Tsunemi et al., 2020; Wong, 2020), understanding the role of BAG3 in these endocytic processes and how it impacts tau uptake and processing is essential. Indeed, recent studies provide evidence that deletion of BAG3 from astrocytes impaired Aβ clearance in co-cultures with APP/PSEN1 neurons, indicating that astrocyte BAG3 plays a role in AD-relevant processes (Augur et al., 2025). Therefore, in this study, we investigated the role of BAG3 in mediating CME and tau uptake and processing in primary astrocytes, beginning with investigation of potential interactors. Clathrin heavy chain was previously identified as a BAG3 interactor in an unbiased screen of BAG3 interactors (Lin et al., 2022b), and via our proximity ligation assay (PLA) in this study, we show that BAG3 and clathrin heavy chain closely associate in astrocytes (<40 nm) (Alam, 2018). We also demonstrate that depleting BAG3 significantly decreases clathrin-AP-2 interactions. Ligand bound epidermal growth factor (EGF)-receptor is predominantly internalized by CME (Narayan et al., 2020; Zhou and Sakurai, 2022), and we show that this process is significantly inhibited by depletion of BAG3. Previously, it was demonstrated that, CME contributes to the uptake of tau monomers by neurons (Evans et al., 2018). Similarly, in astrocytes, we found that the majority of tau uptake was inhibited by a dynamin inhibitor suggesting that CME is involved. Furthermore, depletion of BAG3 also significantly inhibited the initial uptake of tau. Using live cell imaging, we found BAG3 depletion in astrocytes increased the mean lifetime of clathrin particles, indicating an impairment of the assembly, and/or disassembly process of clathrin coated vesicles (CCVs). Additionally, astrocytic BAG3 depletion impacted the endolysosome compartment, demonstrated by an increased number of Lamp1+ puncta, without a change in Lamp1+ puncta size, and an increased co-localization of endocytosed tau with Lamp1+ puncta. In aggregate, the results of these studies clearly indicate that BAG3 plays a pivotal role in mediating CME and tau uptake in astrocytes and that it modulates vacuolar processes in astrocytes. Additionally, BAG3 may also play an important role in regulating the properties of the vacuoles that contain tau and other CME cargo. Given the key role that astrocytes play in mediating the progression of tau pathology (Eisenbaum et al., 2024; Fleeman and Proctor, 2021), these studies indicate that BAG3 may play a protective role in part by facilitating CME and tau clearance in astrocytes and that deficits in BAG3 could contribute to disease pathogenesis.

## METHODS AND MATERIALS

### Virus preparation

Lentiviral particles were prepared using a lentiviral vector with a U6 promoter, pHUUG (a generous gift of from Dr. C.Pröschel) expressing BFP only (empty vector (EV) control) or with shBAG3 (5’-AAGGTTCAGACCATCTTGGAA-3’) (Lin et al., 2022b). HEK cells were transfected with lentiviral packaging vectors and either shBAG3 or EV lentiviral vectors. Lentiviruses were collected as we have described previously (Lin et al., 2024; Lin et al., 2022b). After concentration, the lentiviruses were aliquoted, snap frozen, and stored at -80°C. Virus was thawed on ice immediately prior to use.

### Astrocyte culture

Primary astrocytes were cultured at post-natal day 0 from wild-type C57BL/6 mouse pups as previously described (Delgado et al., 2024; Emerson et al., 2023). In brief, the brains were dissected, meninges were removed, and cortical hemispheres were collected. Following trituration of the cells, they were plated onto culture dishes in MEM supplemented with 10% FBS, 33 mM glucose, 1 mM sodium pyruvate (Gibco, 11360-070), and 0.2% Primocin (Glial MEM). Twenty-four hours later, the dishes were shaken vigorously and rinsed to remove debris and other cell types. Astrocytes were maintained at 37 °C/5% CO_2_ for 7–8 days until near confluency, frozen in Glial MEM containing 10% DMSO, and stored in liquid nitrogen. For all studies astrocytes were thawed and plated onto 60mm plates until they had reached 80%+ confluency. Astrocytes were then trypsinized and split onto coverslips or culture plate dishes as needed for the studies. On day 3 after replating astrocytes, lentivirus was added. Media was changed after 24 hours and incubation continued for 7-8 days with a half media change every 4 days.

### Preparing and labeling recombinant tau

Recombinant tau was purified from E. Coli BL21 (DE3) as previously described (KrishnaKumar and Gupta, 2017) with some modifications. Plasmid PTC7 0N4R (a generous gift from J. Kuret, Ohio State University) was transformed into E. Coli BL21 (DE3) via heat shock. A single colony was picked from a selective agar plate and inoculated in 25mL terrific broth containing 100µg/mL ampicillin and 50µg/mL chloramphenicol. This culture grew overnight at 37°C in a shaker (250 rpm). The next morning, the starter culture was added to 1L terrific broth containing the same antibiotics. When A_600_ reached 0.6-0.8, IPTG (Invitrogen, 15529-019) was added to a final concentration of 1mM to induce tau expression. Cultures were allowed to grow for 4 hours with continuous shaking. Cells were pelleted by centrifugation at 7,500xg for 10 minutes at 4°C. Cell pellets were frozen at -80°C. The next day, the pellets were thawed and resuspended in 40mL lysis buffer (0.5M NaCl, 10mM Tris, pH 8.0) containing protease inhibitors. The suspension was subsequently incubated at 100°C for 20 minutes. After cooling, the suspension was centrifuged at 10,000xg for 45 minutes at 4°C. The supernatant was collected, passed through a 5µm filter, and then dialyzed into PBS overnight at 4°C. The filtrate contained tau was analyzed by SDS-polyacrylamide gel electrophoresis (PAGE) and Coomassie blue staining to measure concentration using BSA as a standard. Once concentration was calculated, appropriate amounts of Dylight-488 NHS ester was added to the solution for 30 minutes. The Dylight 488 labeled tau (488-tau) was again dialyzed into PBS overnight at 4°C; then it was aliquoted and frozen at -80°C. The final 488-tau stock was analyzed by SDS- PAGE with Coomassie blue staining to confirm the final concentration and purity.

### EGF and tau uptake

Astrocytes were incubated for 1 hour at 37°C and 5% CO_2_ in serum-free MEM media prior to any uptake. Following the protocol of Narayan et al. (Narayan et al., 2020), astrocytes were removed from the incubator and placed on ice for 5 minutes to halt endocytic processes. The MEM media was removed and astrocytes were washed 1X with PBS (Corning, 21-031-CM). Then, serum free media with 100ng/mL EGF-488 or 300 nM 488-tau was added and the astrocytes returned to the incubator for the incubation times indicated prior to fixation and analyses. For EGF uptake, only flow analysis was performed.

For immunocytochemistry (ICC) and proximity ligation assay (PLA), media was removed from the astrocytes which were then washed 1X with PBS and fixed in 4% paraformaldehyde (PFA) for 5 minutes. Astrocytes were then washed with 1X PBS, then 0.25% Triton-X was added for 10 minutes to permeabilize the cell’s membranes. Astrocytes were then washed 1X with PBS and blocked in a mixture of 5% BSA in PBS and 1.5 M glycine for 1 hour. Astrocytes were washed again in 1X with PBS and appropriately labeled with primary antibody diluted in 5% BSA. Astrocytes were incubated with primary antibodies were at 4 °C overnight before being washed with 1X PBS. For ICC, fluorescently tagged secondary antibodies were added at a concentration of 1:1000 for 1 hour, followed by being washed in 1X PBS then mounted. Cells were mounted using EMS Shield Mount with DABCO (EMS, 17985-150). Processing of fixed astrocytes for PLA is described below.

For flow cytometry, media was removed, and astrocytes washed 1X with cold PBS. Trypsin (Corning, 22-051-CI) was added, and astrocytes placed back into the CO_2_ incubator for 5 minutes. Trypsin was then neutralized by adding serum containing MEM. Suspended astrocytes were then collected into 15 ml conical tubes and spun for 5 minutes at 500xg. Excess supernatant was removed, and the pellet was resuspended in 2mL of PBS and passed through a 40 µm cell strainer. An equal volume of 4% PFA was added [final concentration = 2%] and allowed to fix for 4 minutes. Cells were then centrifuged at 500xg for 5 minutes. The pellet was resuspended in approximately 75 µL PBS ready for flow cytometry.

### Flow cytometry

Flow cytometry was carried out using an ImageStream GenX with a laser power of 100mW in the URMC Flow Cytometry Resource Core (RRID: SCR_012360). For most analyses, approximately 10,000 astrocytes were counted in each condition. Images were then analyzed using IDEAS software. Images were thresholded to ensure only single cells of a particular size and circularity (aspect ratio: 0.6-1) were analyzed. Appropriate images were analyzed using Spot count wizard in the IDEAS program to generate a spot count for each cell. These spot counts were then statistically analyzed using GraphPad Prism.

### Immunocytochemistry analysis

Confocal images were captured with an Olympus FV1000MP microscope and a UPlanSApo 60x oil immersion objective. Images were analyzed using Fiji ImageJ. Images were thresholded using ImageJ’s threshold tool (images were threshold to only contain pixels of greater than 1044 value), and spots were analyzed using the analyze particle’s function (pixel size 0-4, circularity 0.00-1.00).

### Live cell imaging of clathrin dynamics

Astrocytes were transduced with either shBAG3/BFP or BFP-only (EV) lentiviruses 7 days prior to transfection with a construct expressing mCherry fused to the N terminus of clathrin light chain A (CLC) using Lipofectamine 2000 for 48 hours prior to imaging. mCherry-CLC is a structurally and functionally active fusion protein (Ganguly et al., 2021). Live-imaging was performed using the Teledyne Photometrics Prime BSI express sCMOS camera mounted on the Nikon ECLIPSE Ti2 inverted microscope equipped with the NIS-Elements 6D imaging acquisition module (v.5.42.06). The Nikon D-LEDI fluorescence LED illumination system (equipped with 385 nm, 488 nm, 568 nm and 621 nm excitation wavelengths) was used as the primary illumination source. Specific illumination wavelengths were selected by combining a large field of view quad-filter cube (DAPI/FITC/TRITC/CY5; 96378) with specific Lumencor emission filters (FF01-474/27-32, FF01-515/30-32, FF01-595/31-32, FF02-641/75-32) https://www.nature.com/articles/s41586-025-09018-7#Sec9. Videos were captured using a CFI Plan Apochromat VC 100 Å∼ oil (NA 1.40; Nikon) objective from cell incubated in the Tokai Hit Stage top incubator at 37C in Hibernate-E low fluorescence imaging solution (Ganguly et al., 2021). Astrocytes were selected to ensure they displayed signs of transduction and were efficiently transfected. Movies were captured at 1 frame per second with an exposure of 500ms at 15% LED power for 6 minutes. The resulting video was analyzed using ImageJ’s (FiJi) kymograph builder tool (https://www.nature.com/articles/s41467-025-57135-8).

### Proximity ligation assay

Astrocytes were fixed and permeabilized and subsequently processed using the Duolink proximity ligation assay kit (Sigma, DUO92002-30RXN). After primary antibodies were added, cells were incubated overnight at 4°C. The next day, cells were washed with 1X PBS. Duolink secondary antibodies diluted 1:5 in the Duolink antibody diluent were added to the astrocytes. Cells were incubated at 37°C for 1 hour. Cells were washed with 1X PBS. 5X Duolink ligation buffer was diluted 1:5 in HPLC water, and ligase was added at a concentration of 1:40. Ligation buffer was added, and cells were incubated at 37°C for 30 minutes. Cells were washed with 1X PBS. 5X amplification buffer was diluted 1:5 in HPLC water. Polymerase was added to the diluted amplification buffer at a 1:80 dilution. Polymerase buffer was added to the cells followed by incubation at 37°C for 100 minutes. Cells were then washed and mounted on to a slide (EMS, 17985-150) for confocal imaging. It should be noted that all PLA measures were done after incubation with 300 nM 488-tau for 5 minutes.

### Immunoblotting

Astrocytes were collected using 1X RIPA buffer with a cocktail of protease inhibitors. Cells were sonicated for 15 seconds and centrifuged at 16,000xg for 10 minutes at 4°C. The supernatant was collected and protein concentration measured by BCA. Samples were made up at 1mg/mL in 1X SDS buffer. Prior to loading, samples were boiled for 10 minutes at 100°C. Ten micrograms of protein was run on the gel.

### Reagents

Reagents:
| Reagent | Supplier | Catalog # |
| --- | --- | --- |
| BAG3 primary antibody | Proteintech | 10599-I-AP |
| Beta actin primary antibody | Proteintech | 66009-1-ig |
| Duolink proximity ligation assay kit | Sigma | DUO92002-30RXN,<br>DUO92008-30RXN |
| BL21(DE3) Competent *E. coli* | New England Biolabs | C2527H |
| EMS shield mount with DABCO | EMS | 17985-150 |
| Lipofectamine 2000 | Thermo Fisher Scientific | 11668030 |
| DyLight™ 488 NHS Ester | Thermo Fisher Scientific | 46402 |
| PBS | Corning | 21-031-CM |
| Trypsin | Corning | 22-051-CI |
| GlutaMAX supplement | Gibco | 41090036 |
| Fetal Bovine Serum | Atlas | F-500-DR |
| Primocin | Invivogen | ant-pm-1 |
| Dynasore | Cayman Chemicals | 14062 |
| Dylight 488 NHS ester | Thermo Fisher Scientific | 46403 |
| LAMP1 primary antibody | Santa Cruz | sc-20011 |
| Vimentin primary antibody | Cell Signaling | 5741S |
| Cathepsin D primary antibody | Proteintech | 21327-1-AP |
| Clathrin primary antibody | Proteintech | 10852-1-AP |
| AP-2 primary antibody | Novus | NB600-1545 |
| Alexa Fluor® 488 labeled EGF | Invitrogen | E13345 |
| GFAP primary antibody | Abcam | Ab53554 |

### Statistical Analysis

All statistical analyses were performed using GraphPad Prism 10. All data were subject to an outlier analysis using ROUT (Q=0.5%). Cleaned data was subjected to a Welch’s unpaired T test to identify level of significance.

## RESULTS

To deplete BAG3, astrocytes were transduced with a shBAG3 lentivirus (Lin et al., 2022b) or with a control lentivirus expressing BFP only (empty vector; EV) for 7 days. For all imaging experiments, parallel plates were transduced, collected and immunoblotted to confirm knockdown of BAG3. BAG3 levels were reduced 85.0+/- 6.4% by shBAG3 transduction in all experiments (Figure 1). Previously, using an unbiased proteomic approach we identified a number of CME proteins as BAG3 interactors (Lin et al., 2022b). To extend these findings, we used PLA to examine the association of BAG3 with clathrin. A positive PLA signal indicates that the two proteins are within 40 nm or less of each other, strongly indicating an interaction (Alam, 2018). Images of PLA signal for BAG3-clathrin (heavy chain) are shown in Figure 2A. Quantified data in Figure 2B demonstrate that the number of PLA puncta is significantly decreased when BAG3 is depleted. In addition to clathrin, the unbiased proteomic analyses showed that BAG3 interacted with a number of subunits of the AP-2 complex (Lin et al., 2022b). Since the AP-2 complex plays an essential role in initiating the formation of clathrin coated pits (Zaccai et al., 2022), we next examined the impact of BAG3 depletion on clathrin (light chain)/AP-2 interactions. The representative PLA signaling shown in Figure 2C and the quantified data shown in Figure 2D demonstrates that BAG3 depletion significantly decreases clathrin/AP-2 interactions. Given that BAG3 interacts with key proteins of CME, and that ligand-bound EGF-receptor internalization is often used to measure CME-mediated import (Narayan et al., 2020; Zhou and Sakurai, 2022), we next measured the role of BAG3 in mediating CME in astrocytes using flow cytometry to quantify the uptake of Alexa Fluor® 488-labeled EGF. These measurements were first carried out in the absence or presence of the small molecule dynamin inhibitor, Dynasore (100µM) (Evans et al., 2018; Macia et al., 2006), and clearly demonstrated that inhibition of dynamin inhibits the majority of EGF uptake (Figure S1). We then depleted BAG3 in the astrocytes, followed by measurement of 488-labeled EGF uptake. These data demonstrate that depletion of BAG3 significantly decreased EGF uptake (58.8 +/- 0.1% of control) (Figure 2E). Violin plots showing data as spot count per cell for individual biological replicates are in Figure S2.

**Figure 1.**
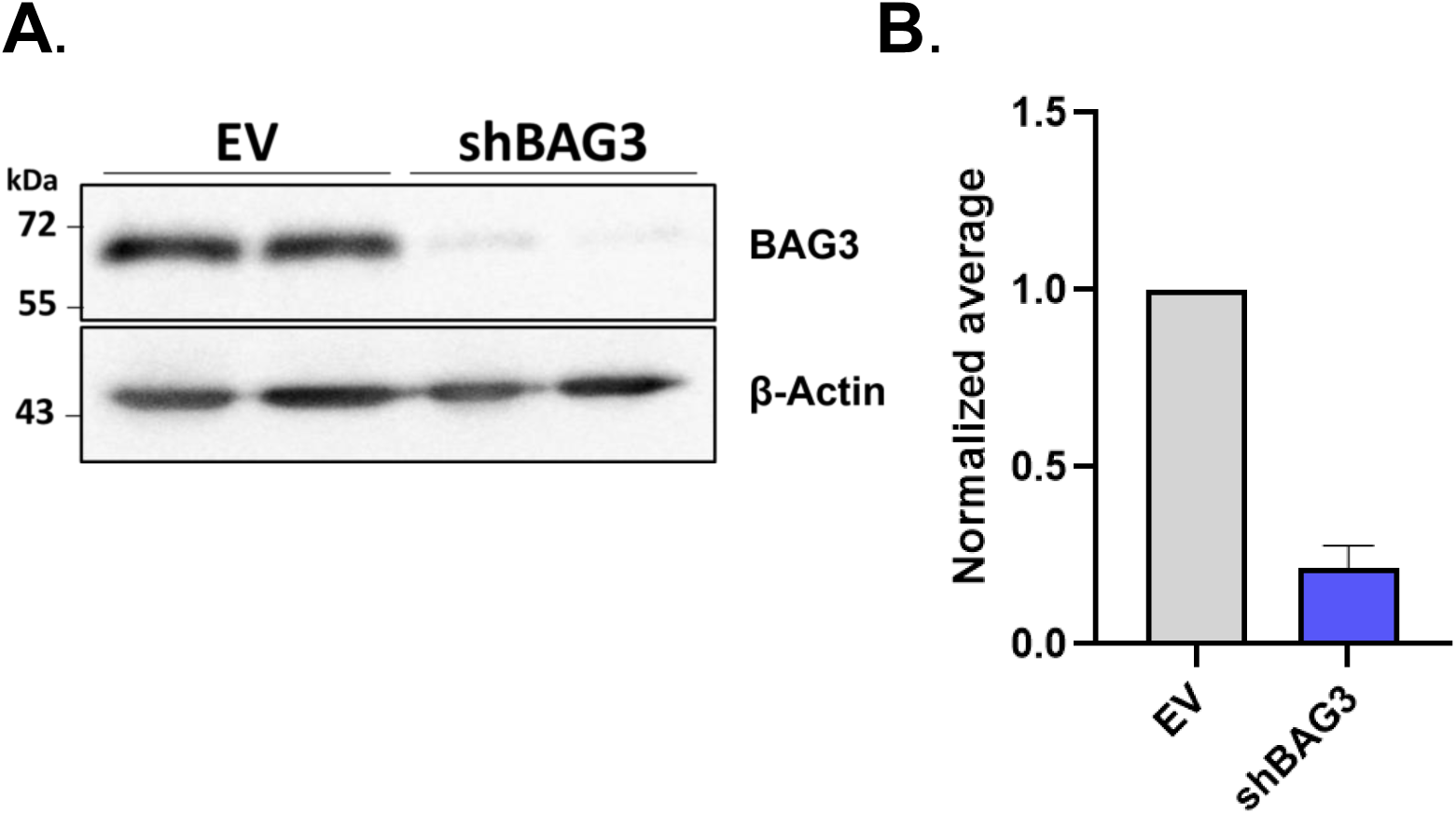
BAG3 is efficiently depleted in astrocytes transduced with shBAG3. (A) Representative immunoblot of astrocytes transduced with lentivirus expressing empty vector (EV) or BAG3 shRNA for 6-7 days. Cell lysates were immunoblotted for BAG3 validating knockdown, and β-actin was used as a loading control. (B) Quantitative analysis of BAG3 immunoblot data demonstrating 85 +/- 6% knockdown in the shBAG3 condition (n=6).

**Figure 2.**
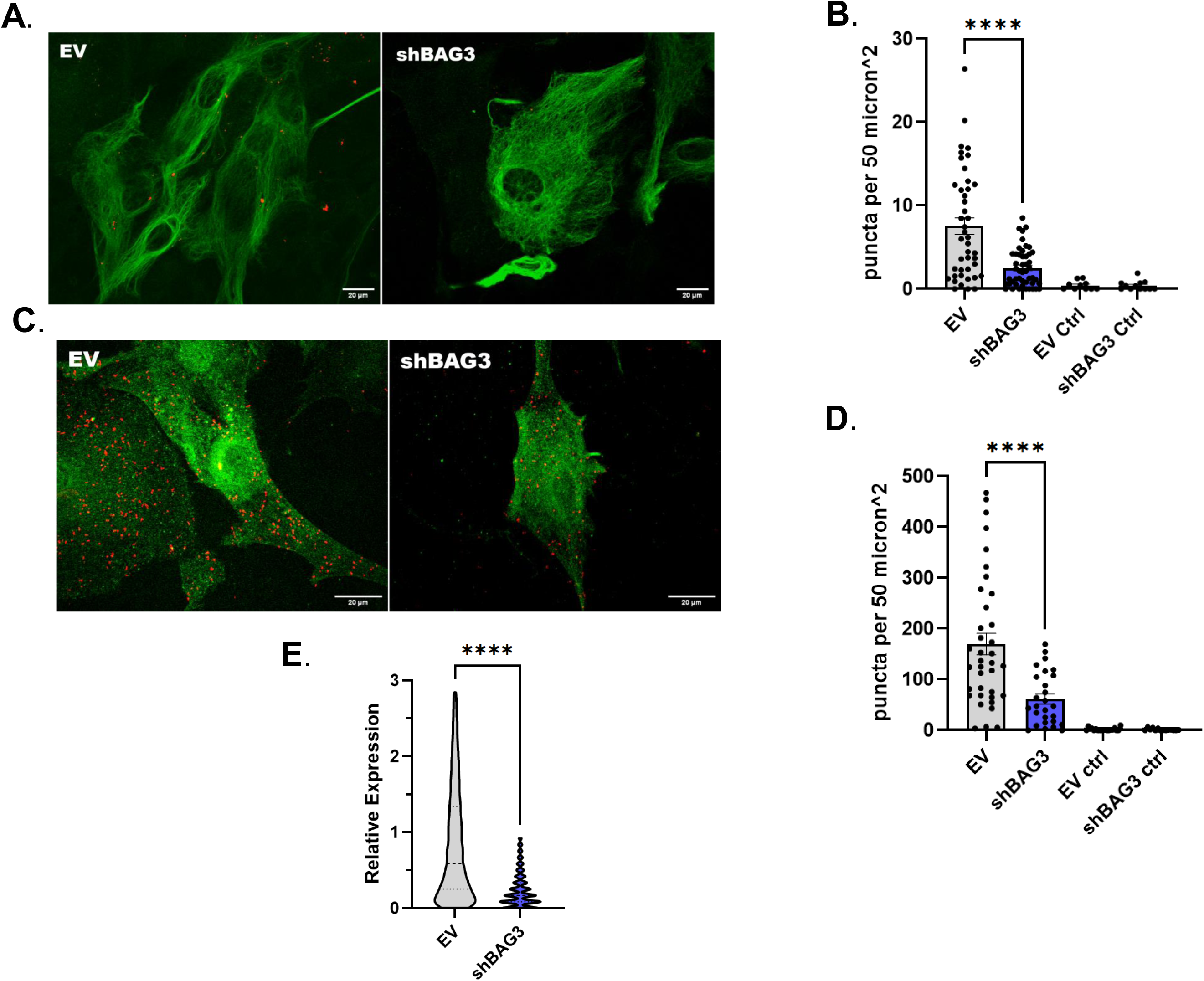
BAG3 interacts with and regulates CME in astrocytes. (A) Representative PLA results showing BAG3 and clathrin interact (red puncta in images) and that depleting BAG3 significantly reduces the PLA signal. Astrocytes were co-stained for GFAP and pseudo colored green. Scale bar = 20µm. (B) Quantification of PLA puncta demonstrating that depletion of BAG3 significantly reduces the number of BAG3 and clathrin interactions as indicated by a reduction in red PLA puncta. Controls were processed identically, except the ligase step was omitted. (****p<0.0001, n=2 biological replicates with >30 cells analyzed for each condition). (C) Representative PLA results showing AP-2 and clathrin interactions are significantly decreased when BAG3 is depleted in astrocytes as indicated by a reduction in the red PLA puncta. Cells were co-stained with GFAP and pseudo colored green. Scale bar = 20µm. (D) Quantification of PLA puncta demonstrating a significant decrease in AP-2 and clathrin interactions in BAG3 depleted astrocytes. Controls were incubated without ligase. (****p<0.0001, n=2 biological replicates with >30 cells analyzed for each condition). (E) Flow cytometry data showing BAG3 depletion significantly inhibits EGF uptake by astrocytes. Violin plot shows the relative EGF-488 puncta/cell in the shBAG3 condition compared to EV. Astrocytes were incubated with 100ng/mL EGF-488 for 5 minutes prior to analysis. Raw data was transferred to Graphpad Prism for statistical analysis. Dashed lines represent median values, dotted lines show upper and lower quartiles. (****p <0.0001, n=2 biological replicates, ∼10,000 cells were counted for each replicate and condition). The mean relative value +/- SD plotted on the violin plot for EV was 0.831 +/-0.731 and for shBAG3 it was 0.233 +/- 0.228. The median of the relative value for EV was 0.584 and for shBAG3 it was 0.167.

Previous studies have provided evidence that neurons take up monomeric tau through CME (Evans et al., 2018). To determine if astrocytes also use CME to take up tau, astrocytes were incubated with 300 nM 488-tau for 5 min in the absence or presence of Dynasore prior to collection and flow cytometry. These data demonstrate that a dynamin-dependent process contributes to tau uptake tau by astrocytes (50.5 +/- 0.1% of control) (Figure 3A). Violin plots showing data as spot count per cell for individual experiments are in Figure S3. We next determined how BAG3 depletion in astrocytes impacted tau uptake. These data demonstrate that BAG3 depletion significantly decreases tau uptake by astrocytes (77.4 +/- 0.1% of control) (Figure 3B), which is similar to the inhibition observed when measuring EGF uptake, which is predominantly CME dependent. Violin plots showing data as spot count per cell for individual experiments are in Figure S4.

**Figure 3.**
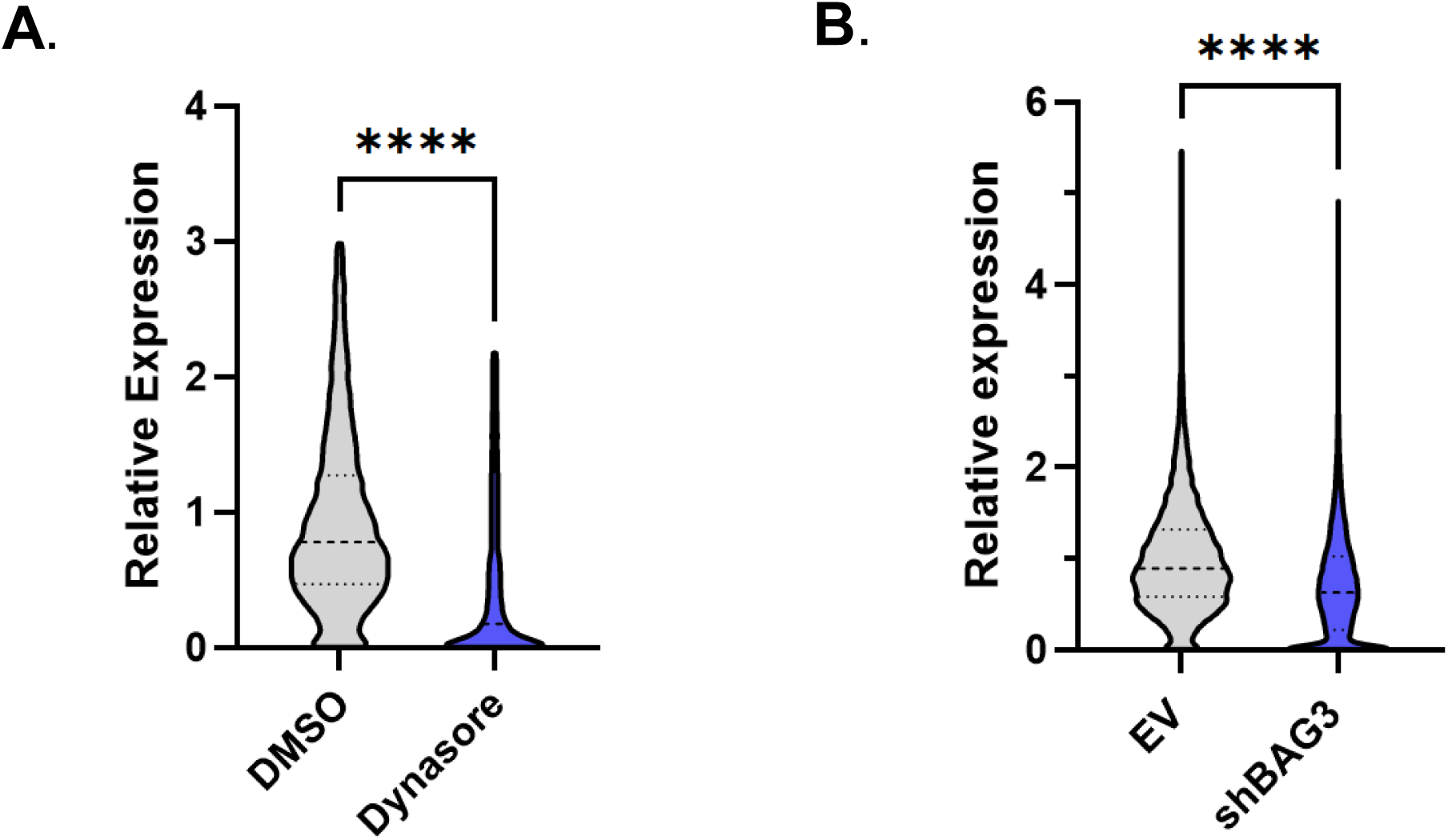
Tau is endocytosed by astrocytes via a dynamin/CME dependent process which is inhibited by depletion of BAG3. Cells were incubated with 300 nM 488-tau for 5 minutes. Raw data was transferred to Graphpad Prism for all statistical analyses. Dashed lines on violin plots represent median values, dotted lines indicate an upper and lower quartile. (A) Flow cytometry data showing inhibition of dynamin with Dynasore (100 µM) significantly inhibits uptake of tau by astrocytes. Violin plot demonstrating relative number of 488-tau puncta/cell in the presence of Dynasore compared to DMSO control. Cells were pre incubated with 100µM Dynasore or DMSO for 30 minutes prior to the addition of 300 nM 488-tau. (****p <0.0001, n=2 biological replicates, ∼10,000 cells were counted for each replicate and condition). The mean relative value +/- SD plotted on the violin plot for DMSO was 0.929 +/- 0.651 and for Dynasore it was 0.470 +/- 0.601. The median of the relative value for DMSO was 0.778 and for Dynasore was 0.174. (B) Flow cytometry data showing depletion of BAG3 significantly inhibits uptake of tau by astrocytes. Violin plot shows relative value of 488-tau puncta per cell in the shBAG3 condition relative to empty vector (EV). (****p <0.0001, n=3 biological replicates, ∼10,000 cells were counted for each replicate and condition). The mean relative value +/- SD plotted on the violin plot for EV was 0.999 +/- 0.598 and for shBAG3 it was 0.693 +/- 0.583. The median relative value for EV was 0.895 and for shBAG3 was 0.628.

Next, we wanted to determine how BAG3 depletion in astrocytes impacted the dynamics of clathrin particles. Astrocytes were transduced with shBAG3 or control lentivirus, then, after 6 days, they were transfected with mCherry-CLC. Forty-eight hours later, they were imaged every second for 6 minutes. Representative kymographs are shown in Figure 4A; red arrowheads indicate abrupt disappearance of fluorescence indicating disassembly (Ganguly et al., 2021). Quantification of the average lifetime of the clathrin particles (Figure 4B) demonstrate that BAG3 depletion significantly increases the average clathrin particle lifetime (Ganguly et al., 2021) suggesting a slowing of CME kinetics. Data is presented as both raw data seconds and relative lifetime.

**Figure 4.**
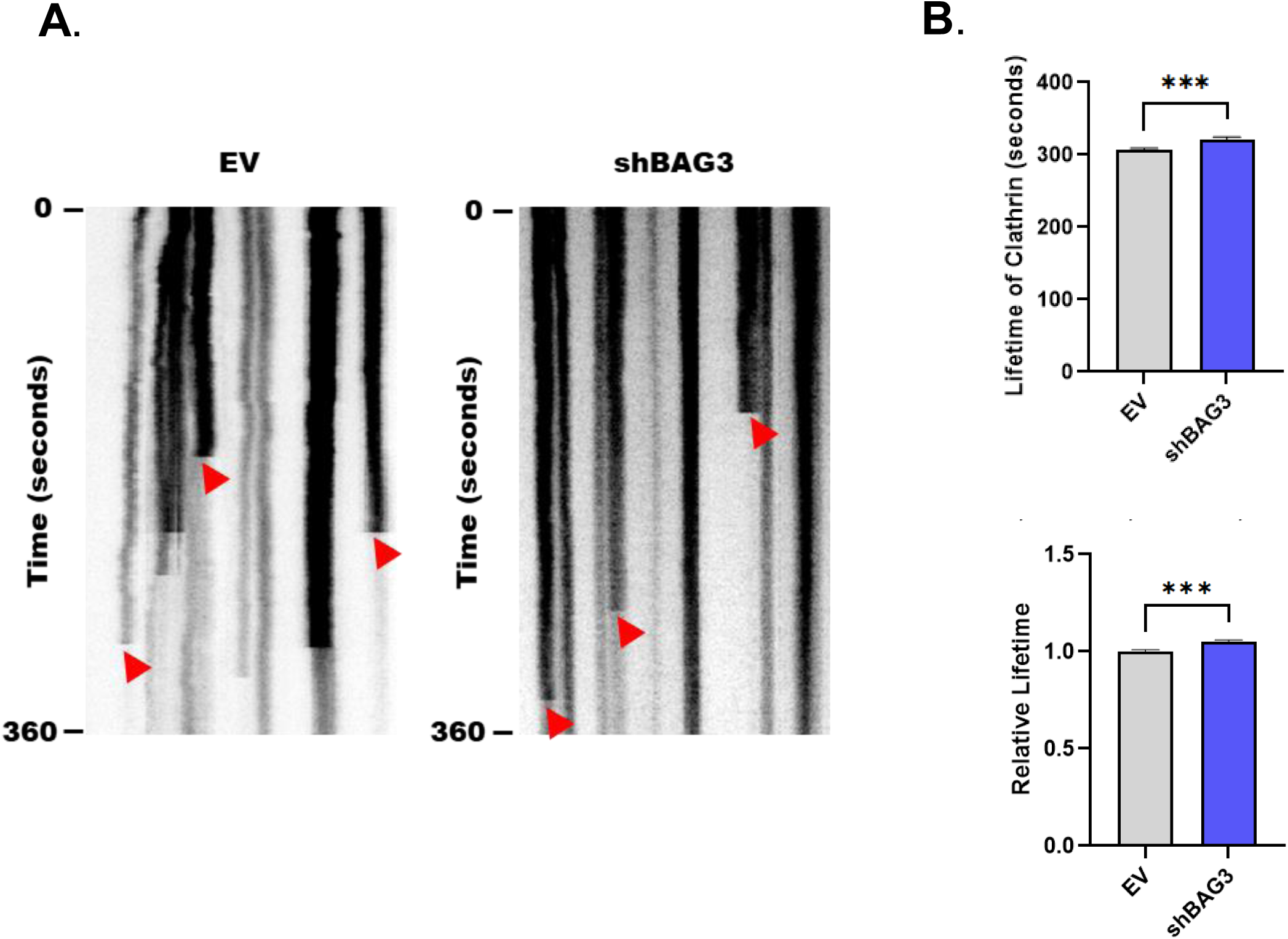
BAG3 depletion in astrocytes significantly increases mCherry-CLC lifetime. (A) Representative kymographs of mCherry-CLC particle lifetimes in EV and shBAG3 conditions. Y axis is time in seconds. Kymographs were produced using ImageJ’s KymographBuilder plug in. Red arrowheads indicate representative points where there is a sudden loss of mCherry-CLC signal and where the lifetime of a particular puncta would be measured. (B) Quantitative analysis of the lifetime of mCherry-CLC particles in EV and shBAG3 transduced astrocytes. ImageJ was utilized to measure lifetime of each Clathrin light chain particle, and raw data was transferred to Graphpad Prism for statistical analysis. Top graph shows raw data and bottom graph shows data expressed as relative life time. ***p< 0.001, n=2 biological replicates, 7 cells were analyzed for each replicate and condition.

In CME, once the clathrin-coated vesicles are formed and released from the membrane they uncoat, and the endosomes progress through the endocytic pathway (Mettlen et al., 2018). Previous studies demonstrated that tau taken up by neurons, predominantly by CME, was trafficked to Lamp1+ endolysosomes (Evans et al., 2018). Therefore, astrocytes with and without depletion of BAG3 were incubated for 45 minutes in the presence of 488-tau prior to measurement of Lamp1+ structures. Figure 5A shows representative images of Lamp1 and 488-tau punctae in the astrocytes. Quantification of the number of Lamp1+ punctae showed a significant increase in shBAG3-transduced astrocytes compared to control (Figure 5B), with no effect on the average size of the puncta (Figure 5C). However, depleting BAG3 in astrocytes resulted in increased co-localization/co-occurrence of 488-tau with Lamp1 puncta using Manders coefficient (Aaron et al., 2018) (Figure 5D).

**Figure 5.**
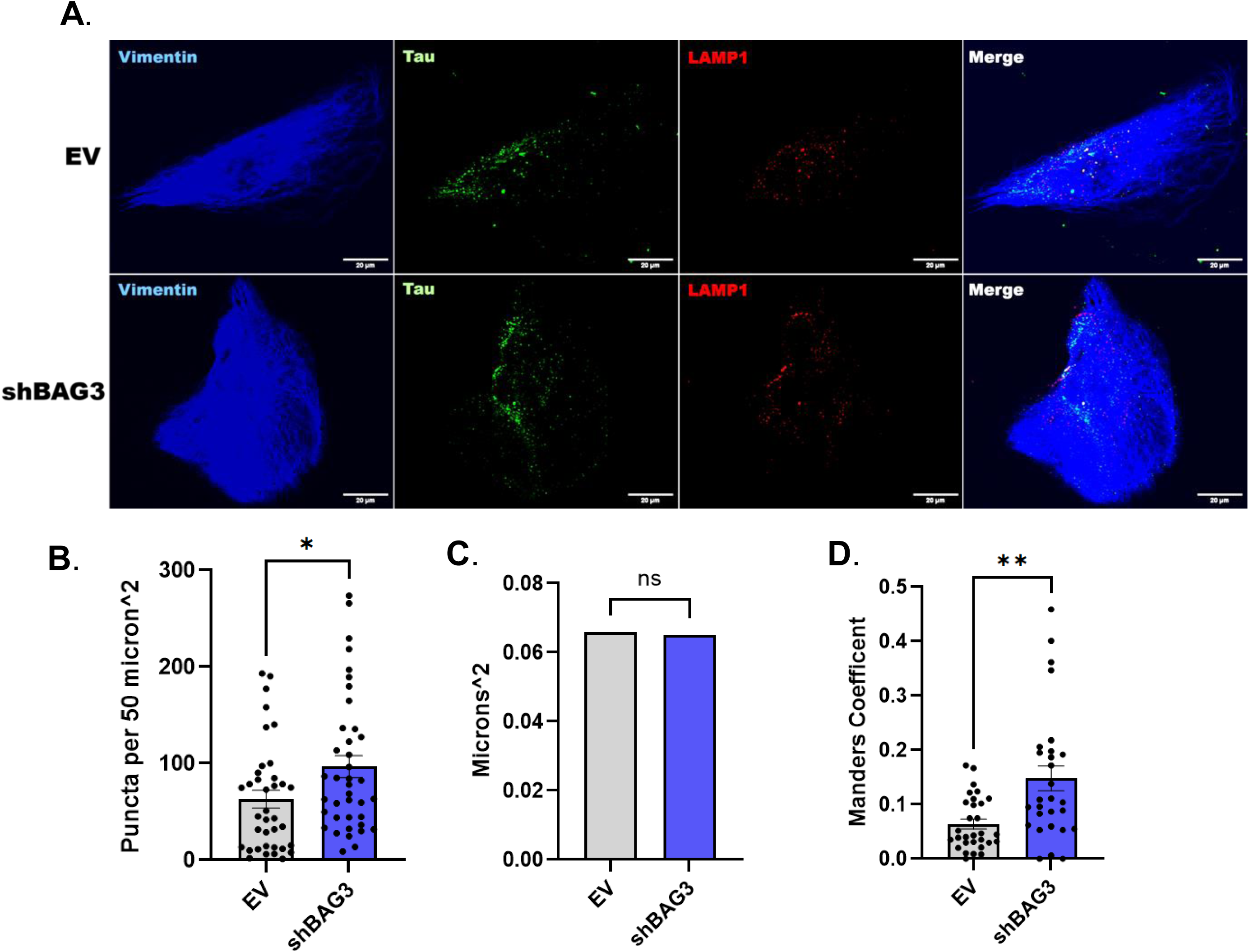
BAG3 depletion in astrocytes significantly increases the number but not the size of Lamp1 compartments, and the co-localization of endocytosed 488-tau with Lamp1 puncta. (A) Representative images of EV and shBAG3 transduced astrocytes after 45 minute incubation with 300 nM 488-tau (green puncta). Astrocytes were stained for vimentin (blue) to identify the cells and a LAMP1 antibody (red puncta). Scale bar = 20µm. (B) Quantification of number of LAMP1+ puncta present in the EV and shBAG3 conditions. Images were thresholded and analyzed using ImageJ’s analyze particles function (pixel size 0-4, circularity 0.00-1.00). Each puncta was counted as its own data point. (*p<0.05, n=3 biological replicates, 10 cells for each condition and replicate were analyzed, 8512 total puncta analyzed for EV, 8353 total puncta for shBAG3). (C) Quantification of the size of LAMP1 positive vacuoles present in EV and shBAG3 conditions. Images were thresholded and analyzed using ImageJ’s analyze particles function (pixel size 0-4, circularity 0.00-1.00). (P value = 0.1430, N.S., n=3 biological replicates, 10 cells for each condition and replicate were analyzed). D) Mander’s Coefficient of 488-tau and LAMP1 co-localization in the EV and shBAG3 conditions. Images were thresholded and analyzed using ImageJ’s JACoP plugin (threshold = 1044). **p<0.01, n=3 biological replicates, 10 cells for each condition and replicate were analyzed.

## DISCUSSION

Previous studies have provided evidence that BAG3 plays a key role in mediating cellular proteostasis in general (Behl, 2016; Sturner and Behl, 2017) and, more specifically, vacuolar dependent processes (Lin et al., 2023; Lin et al., 2022a). Using neuronal model systems, studies have shown that BAG3 is an important regulator of autophagy (Gamerdinger et al., 2009; Lei et al., 2015; Lin et al., 2024) and Endosomal Sorting Complexes Required for Transport (ESCRT)/Endolysosome pathway function (Lin et al., 2022a). BAG3 is likely involved in neurodegenerative conditions characterized by the accumulation of aggregate prone proteins. BAG3 levels are significantly higher in inhibitory neurons, which are resistant to tau pathology compared to excitatory neurons, which are more susceptible (Fu et al., 2019). Furthermore, BAG3 expression decreases the accumulation of hyperphosphorylated tau in AD models (Sweeney et al., 2024). Deletion of BAG3 in neural cells results in a significant increase in tau and phosphorylated tau in mouse models (Lin et al., 2024). BAG3 has also been reported to mediate the autophagic clearance of the Parkinson’s disease (PD)-relevant protein α-synuclein (Cao et al., 2017). Furthermore, recent studies provided evidence that BAG3 is a contributing factor in regulating clearance mechanisms for intrastriatally injected fibrils of tau and α-synuclein by astrocytes; however, the mechanisms involved were not delineated (Sheehan et al., 2023). This finding is intriguing because astrocytes likely play a key role in protecting against the propagation of tau pathology by taking up extracellular tau. However, if the astrocytes are then unable to properly process the tau, it can result in tau-induced neurotoxic astrocytic phenotypes (Martini-Stoica et al., 2018; Reid et al., 2020; Richetin et al., 2020; Sidoryk-Wegrzynowicz et al., 2017). Another study provided evidence that deletion of BAG3 from astrocytes impaired their ability to clear Aβ when co-cultured with neurons from APP/PSEN1 mice (Augur et al., 2025). Additionally, a recent study demonstrated that in AD, reactive astrocytes exhibit a marked decrease in homeostatic genes, which likely results in a loss of their ability to maintain proteostasis and support neuronal health (Dai et al., 2023). These and other findings clearly indicate that inefficient processing of tau and other disease-relevant proteins by astrocytes likely have pathological consequences. Given the function of BAG3 as a mediator of vacuolar clearance processes, it is of critical importance to determine how it functions in astrocytes.

In a previous study, unbiased mass spectrometric analyses revealed that numerous proteins involved in CME were likely BAG3 interactors (Lin et al., 2022a). Here, for the first time we provide evidence that in astrocytes, BAG3 plays a pivotal role in regulating CME. We found that BAG3 closely interacts with clathrin heavy chain, and facilitates AP-2/clathrin interactions, which is essential for the initiation and formation of clathrin-coated pits (Mettlen et al., 2018). Depletion of BAG3 from astrocytes significantly reduced EGF uptake, which was used as a measure of CME (Narayan et al., 2020; Zhou and Sakurai, 2022). These findings suggest that BAG3 may play a supportive role in CME by enabling of AP-2/clathrin interactions. In addition, similar to what was observed with EGF uptake when BAG3 was depleted, tau uptake was also significantly reduced with BAG3 depletion. Previous studies also demonstrated that tau is taken up by neurons through CME (Evans et al., 2018). The fact that treatment with the dynamin inhibitor significantly reduced tau uptake by astrocytes indicates that CME could be a route of entry for tau into astrocytes as it is for neurons. Live cell imaging also supported a role for BAG3 in CME, as depletion of BAG3 in astrocytes increased the lifetime of formed clathrin particles, a finding that likely indicates a decreased rate of CCV uncoating and recycling (Nandez et al., 2014). Once CCVs are uncoated, they fuse with early endosomes and mature into late endosomes which can eventually fuse with lysosomes (McMahon and Boucrot, 2011). In this study we found that there were significantly more endocytosed 488-tau puncta localized to Lamp1+ structures in astrocytes with BAG3 depletion compared to control astrocytes. Previous studies have shown that a significant proportion of LAMP1+ structures do not contain lysosomal hydrolases and are, therefore, likely non-degradative late endosomes that can go on to mature into lysosomes (Cheng et al., 2018). These data suggest that BAG3 maybe facilitating not only CME, but the progression of early endosomes to late endosomes and lysosomes, which is not surprising given that BAG3 is known to interact with several regulators of small GTPases involved in this process. For example,BAG3 interacts with Gapvd1 (also known as Gapex5) (Lin et al., 2022a), a guanine nucleotide exchange factor (GEF) for Rab5 (Kim et al., 2019), which is a key regulator of endosome maturation (Agola et al., 2011; Zhang et al., 2022). BAG3 also interacts with IQ motif and Sec7 domain 1 (Iqsec1), also known as Brefeldin A-resistant ArfGEF 2 (BRAG2) (Lin et al., 2022a), which is a GEF for Arf6 (Fukaya et al., 2020), a primary modulator of intracellular vesicle trafficking (Van Acker et al., 2019). Based on these interactions, it can be speculated that BAG3 depletion initially impairs the rate of cargo uptake by CME; however, due to an impairment and/or delay in endosome maturation and vesicle transport processes, there is an accumulation of cargo (tau) in non-degradative compartments. The impairment of tau processing that results from these alterations in vacuolar dynamics would increase the tau burden within the brain which is a crux of AD pathogenesis. Indeed it can be speculated that an increase in the time that endocytosed tau remains in Lamp1+ organelles could contribute to the formation of tau fibrils/seeds that eventually escape into the cytosol (Chen et al., 2019; Michel et al., 2014). Therefore, this continued thread of investigation has value in establishing potential mechanisms by which compromise of BAG3 function is linked to dementia.

## Supplemental Figures

**Figure S1.**
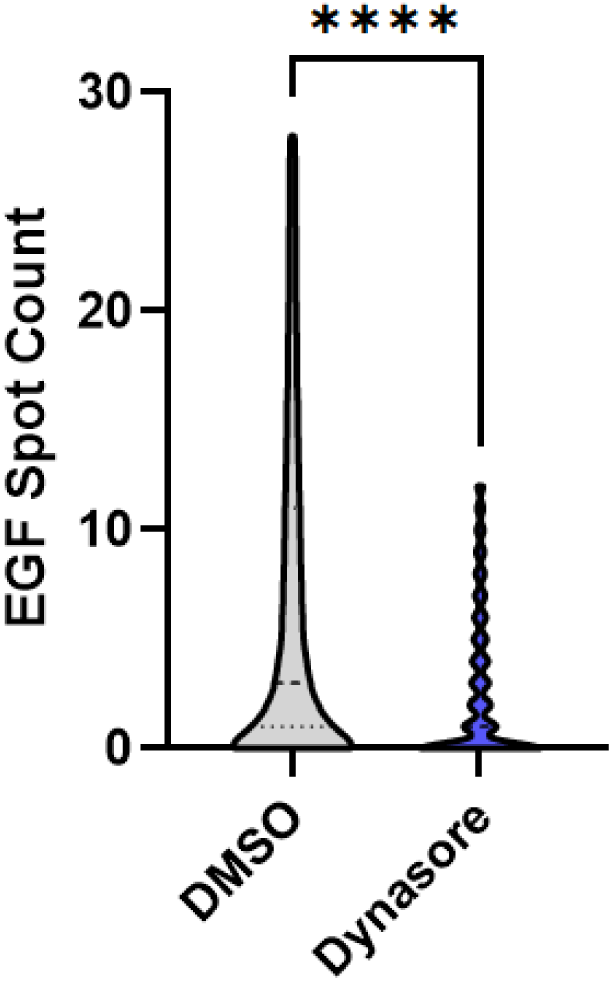
Dynasore significantly inhibits EGF uptake by astrocytes. Astrocytes were pre-incubated with 100µM Dynasore or DMSO for 30 minutes prior to the addition of 100ng/mL EGF-488 for 5 minutes and analysis by flow cytometry. At least 10,000 cells were counted for each group. ****p<0.0001. The mean value +/- SD plotted on the violin plot for EV was 6.530 +/- 7.543 and for shBAG3 it was 0.2.691 +/- 3.421. The median value (dashed lines) for EV was 3.000 and for shBAG3 was 1.000. Dotted lines show upper and lower quartiles.

**Figure S2.**
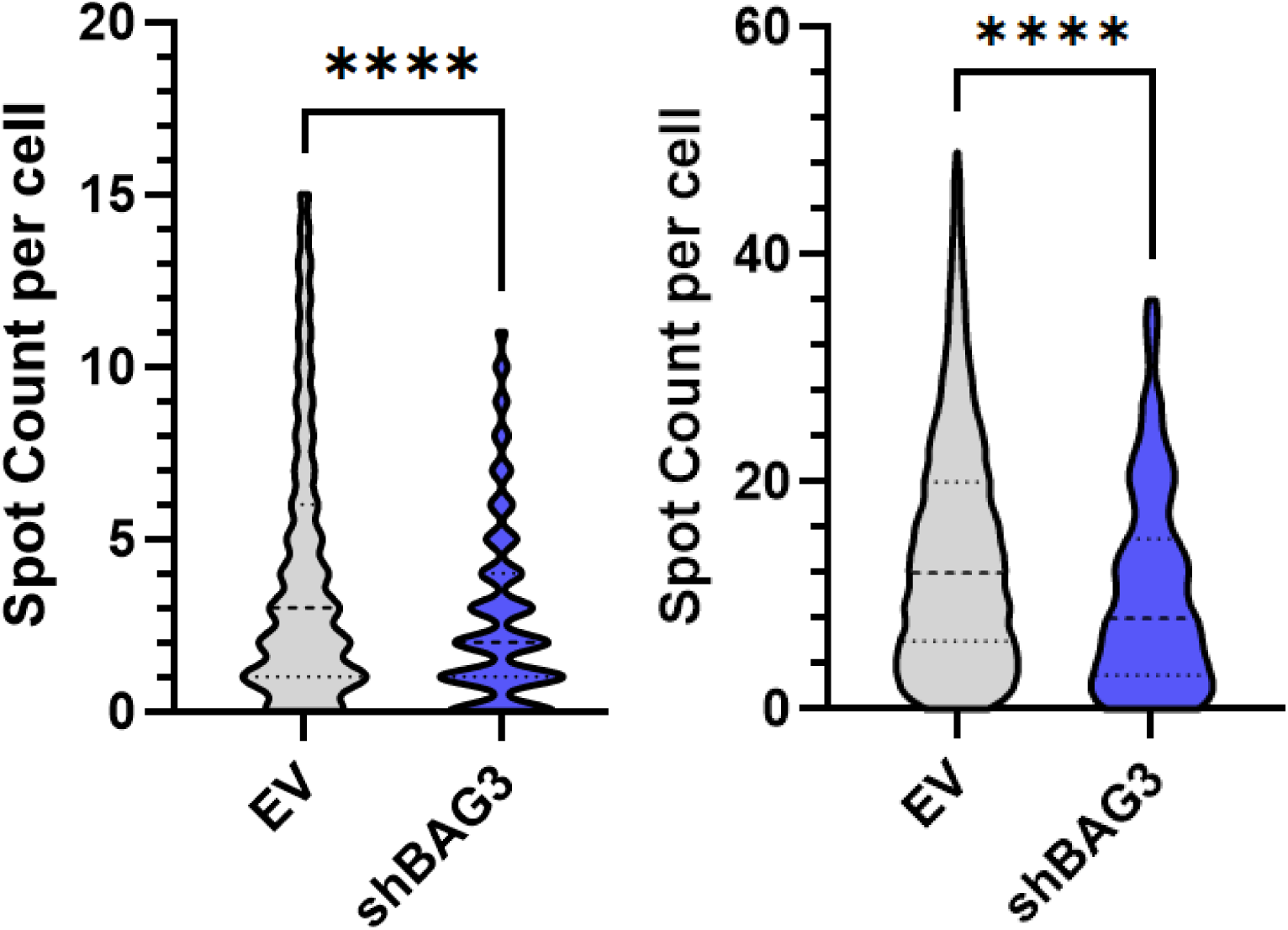
Flow cytometry data presented as violin plots of EGF uptake by astrocytes transduced with EV or shBAG3 as spot count per cell for two individual biological replicates. Individual biological replicates are in each graph. (A) The mean value +/- SD for EV was 3.892 +/- 3.816 and for shBAG3 was 2.750 +/- 2.697. Dashed lines show median values (3.00 for EV and 2.00 for shBAG3) and dotted lines shown upper and lower quartiles. ∼5,000 cells were counted for each group. ****p<0.0001. (B) The mean value +/- SD for EV was 13.99 +/- 10.38 and for shBAG3 was 10.08 +/- 8.628. Dashed lines show median values (12.00 for EV and 8.00 for shBAG3) and dotted lines shown upper and lower quartiles. -10,000 cells were counted for each group. ****p<0.0001. Data analyzed as relative expression are presented in Figure 2E.

**Figure S3.**
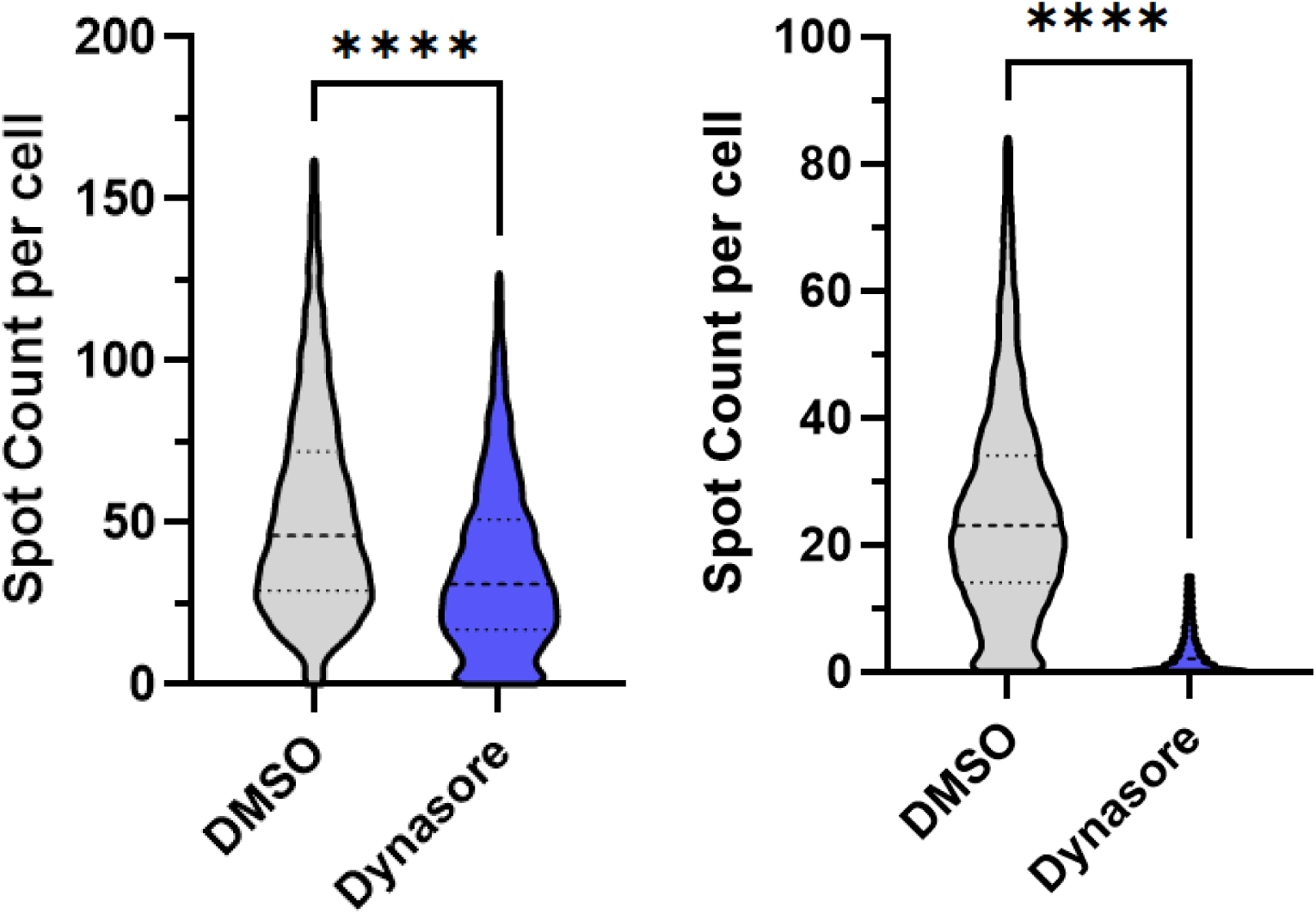
Flow cytometry data presented as violin plots of 488-tau uptake by astrocytes treated with DMSO (control) or Dynasore as spot count per cell for two individual biological replicates. (A) The mean value +/-SD for EV was 53.12 +/- 32.12 and for shBAG3 was 35.97 +/- 25.70. Dashed lines show median values (46.00 for EV and 31.00 for shBAG3) and dotted lines shown upper and lower quartiles. ∼10,000 cells were counted for each group. ****p<0.0001. (B) The mean value for EV was 25.24 +/- 16.74 and for shBAG3 was 3.19 +/- 3.99. Dashed lines show median values (23.00 for EV and 2.00 for shBAG3) and dotted lines shown upper and lower quartiles. ∼10,000 cells were counted for each group. ****p<0.0001. Data analyzed as relative expression are presented in Figure 3A.

**Figure S4.**
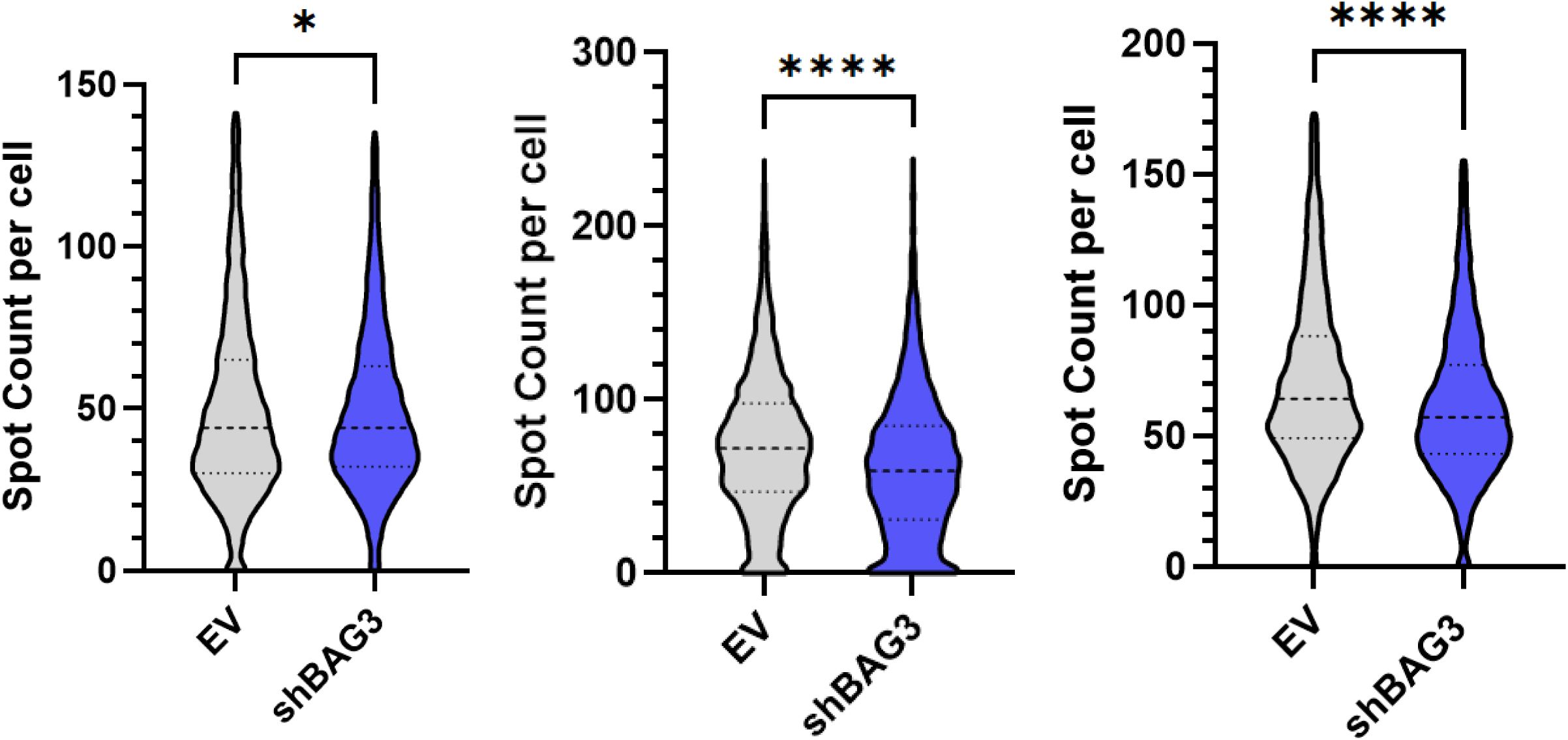
Flow cytometry data presented as violin plots of 488-tau uptake by astrocytes transduced with EV or shBAG3 as spot count per cell for three individual biological replicates. (A) The mean value +/- SD for EV was 49.94 +/- 28.33 and for shBAG3 was 49.18 +/- 25.31. Dashed lines show median values (44.00 for EV and 44.00 for shBAG3) and dotted lines shown upper and lower quartiles. ∼10,000 cells were counted for each group. *p=0.0341. (B) The mean value +/-SD for EV was 73.86 +/- 41.14 and for shBAG3 was 61.28 +/- 40.65. Dashed lines show median values (72.00 for EV and 59.00 for shBAG3) and dotted lines shown upper and lower quartiles. ∼10,000 cells were counted for each group. ****p<0.0001. (C) The mean value +/- for EV was 70.85 +/- 32.32 and for shBAG3 was 62.09 +/- 28.48. Dashed lines show median values (64.00 for EV and 57.00 for shBAG3) and dotted lines shown upper and lower quartiles. ∼10,000 cells were counted for each group. ****p<0.0001. Data analyzed as relative expression are presented in Figure 3B.

## REFERENCES

Aaron, J.S., A.B. Taylor, and T.L. Chew. 2018. Image co-localization - co-occurrence versus correlation. J Cell Sci. 131.

Agola, J.O., P.A. Jim, H.H. Ward, S. Basuray, and A. Wandinger-Ness. 2011. Rab GTPases as regulators of endocytosis, targets of disease and therapeutic opportunities. Clin Genet. 80:305–318.

Alam, M.S. 2018. Proximity Ligation Assay (PLA). Curr Protoc Immunol. 123:e58.

Ando, K., S. Houben, M. Homa, M.A. de Fisenne, M.C. Potier, C. Erneux, J.P. Brion, and K. Leroy. 2020. Alzheimer’s Disease: Tau Pathology and Dysfunction of Endocytosis. Frontiers in molecular neuroscience. 13:583755.

Augur, Z.M., G.M. Fogo, C.R. Benoit, G. Terzioglu, Z.R. Murphy, M.R. Arbery, N. Comandante-Lou, D.M. Duong, N.T. Seyfried, P.L. De Jager, and T.L. Young-Pearse. 2025. BAG3 coordinates astrocytic proteostasis of Alzheimer’s disease-linked proteins via proteasome, autophagy, and retromer complex interactions. bioRxiv:2025.2009.2020.677505.

Behl, C. 2016. Breaking BAG: The Co-Chaperone BAG3 in Health and Disease. Trends in pharmacological sciences. 37:672–688.

Cao, Y.L., Y.P. Yang, C.J. Mao, X.Q. Zhang, C.T. Wang, J. Yang, D.J. Lv, F. Wang, L.F. Hu, and C.F. Liu. 2017. A role of BAG3 in regulating SNCA/alpha-synuclein clearance via selective macroautophagy. Neurobiology of aging. 60:104–115.

Cataldo, A.M., C.M. Peterhoff, J.C. Troncoso, T. Gomez-Isla, B.T. Hyman, and R.A. Nixon. 2000. Endocytic pathway abnormalities precede amyloid beta deposition in sporadic Alzheimer’s disease and Down syndrome: differential effects of APOE genotype and presenilin mutations. Am J Pathol. 157:277–286.

Chen, J.J., D.L. Nathaniel, P. Raghavan, M. Nelson, R. Tian, E. Tse, J.Y. Hong, S.K. See, S.A. Mok, M.Y. Hein, D.R. Southworth, L.T. Grinberg, J.E. Gestwicki, M.D. Leonetti, and M. Kampmann. 2019. Compromised function of the ESCRT pathway promotes endolysosomal escape of tau seeds and propagation of tau aggregation. J Biol Chem. 294:18952–18966.

Cheng, X.T., Y.X. Xie, B. Zhou, N. Huang, T. Farfel-Becker, and Z.H. Sheng. 2018. Characterization of LAMP1-labeled nondegradative lysosomal and endocytic compartments in neurons. J Cell Biol. 217:3127–3139.

Colacurcio, D.J., A. Pensalfini, Y. Jiang, and R.A. Nixon. 2018. Dysfunction of autophagy and endosomal-lysosomal pathways: Roles in pathogenesis of Down syndrome and Alzheimer’s Disease. Free radical biology & medicine. 114:40–51.

Dai, D.L., M. Li, and E.B. Lee. 2023. Human Alzheimer’s disease reactive astrocytes exhibit a loss of homeostastic gene expression. Acta neuropathologica communications. 11:127.

Delgado, T., J. Emerson, M. Hong, J.W. Keillor, and G.V.W. Johnson. 2024. Pharmacological Inhibition of Astrocytic Transglutaminase 2 Facilitates the Expression of a Neurosupportive Astrocyte Reactive Phenotype in Association with Increased Histone Acetylation. Biomolecules. 14.

Eisenbaum, M., A. Pearson, C. Ortiz, M. Mullan, F. Crawford, J. Ojo, and C. Bachmeier. 2024. ApoE4 expression disrupts tau uptake, trafficking, and clearance in astrocytes. Glia. 72:184–205.

Emerson, J., T. Delgado, P. Girardi, and G.V.W. Johnson. 2023. Deletion of Transglutaminase 2 from Mouse Astrocytes Significantly Improves Their Ability to Promote Neurite Outgrowth on an Inhibitory Matrix. Int J Mol Sci. 24.

Evans, L.D., T. Wassmer, G. Fraser, J. Smith, M. Perkinton, A. Billinton, and F.J. Livesey. 2018. Extracellular Monomeric and Aggregated Tau Efficiently Enter Human Neurons through Overlapping but Distinct Pathways. Cell reports. 22:3612–3624.

Fleeman, R.M., and E.A. Proctor. 2021. Astrocytic Propagation of Tau in the Context of Alzheimer’s Disease. Frontiers in cellular neuroscience. 15:645233.

Fu, H., A. Possenti, R. Freer, Y. Nakano, N.C. Hernandez Villegas, M. Tang, P.V.M. Cauhy, B.A. Lassus, S. Chen, S.L. Fowler, H.Y. Figueroa, E.D. Huey, G.V.W. Johnson, M. Vendruscolo, and K.E. Duff. 2019. A tau homeostasis signature is linked with the cellular and regional vulnerability of excitatory neurons to tau pathology. Nature neuroscience. 22:47–56.

Fukaya, M., T. Sugawara, Y. Hara, M. Itakura, M. Watanabe, and H. Sakagami. 2020. BRAG2a Mediates mGluR-Dependent AMPA Receptor Internalization at Excitatory Postsynapses through the Interaction with PSD-95 and Endophilin 3. The Journal of neuroscience : the official journal of the Society for Neuroscience. 40:4277–4296.

Gamerdinger, M., P. Hajieva, A.M. Kaya, U. Wolfrum, F.U. Hartl, and C. Behl. 2009. Protein quality control during aging involves recruitment of the macroautophagy pathway by BAG3. The EMBO journal. 28:889–901.

Ganguly, A., R. Sharma, N.P. Boyer, F. Wernert, S. Phan, D. Boassa, L. Parra, U. Das, G. Caillol, X. Han, J.R. Yates, 3rd, M.H. Ellisman, C. Leterrier, and S. Roy. 2021. Clathrin packets move in slow axonal transport and deliver functional payloads to synapses. Neuron. 109:2884-2901 e2887.

Ginsberg, S.D., M.J. Alldred, S.E. Counts, A.M. Cataldo, R.L. Neve, Y. Jiang, J. Wuu, M.V. Chao, E.J. Mufson, R.A. Nixon, and S. Che. 2010. Microarray analysis of hippocampal CA1 neurons implicates early endosomal dysfunction during Alzheimer’s disease progression. Biol Psychiatry. 68:885–893.

Jaye, S., U.S. Sandau, and J.A. Saugstad. 2024. Clathrin mediated endocytosis in Alzheimer’s disease: cell type specific involvement in amyloid beta pathology. Front Aging Neurosci. 16:1378576.

Kim, H., H. Oh, Y.S. Oh, J. Bae, N.H. Hong, S.J. Park, S. Ahn, M. Lee, S. Rhee, S.H. Lee, Y. Jun, S.H. Kim, Y.H. Huh, and W.K. Song. 2019. SPIN90, an adaptor protein, alters the proximity between Rab5 and Gapex5 and facilitates Rab5 activation during EGF endocytosis. Exp Mol Med. 51:1–14.

KrishnaKumar, V.G., and S. Gupta. 2017. Simplified method to obtain enhanced expression of tau protein from E. coli and one-step purification by direct boiling. Preparative biochemistry & biotechnology. 47:530–538.

Lei, Z., C. Brizzee, and G.V. Johnson. 2015. BAG3 facilitates the clearance of endogenous tau in primary neurons. Neurobiology of aging. 36:241–248.

Lin, H., C.A. Deaton, and G.V.W. Johnson. 2023. Commentary: BAG3 as a Mediator of Endosome Function and Tau Clearance. Neuroscience. 518:4–9.

Lin, H., S.A. Koren, G. Cvetojevic, P. Girardi, and G.V.W. Johnson. 2022a. The role of BAG3 in health and disease: A "Magic BAG of Tricks". Journal of cellular biochemistry. 123:4–21.

Lin, H., S. Sandkuhler, C. Dunlea, J. Rodwell-Bullock, D.H. King, and G.V.W. Johnson. 2024. BAG3 regulates the specificity of the recognition of specific MAPT species by NBR1 and SQSTM1. Autophagy. 20:577–589.

Lin, H., M. Tang, C. Ji, P. Girardi, G. Cvetojevic, D. Chen, S.A. Koren, and G.V.W. Johnson. 2022b. BAG3 Regulation of RAB35 Mediates the Endosomal Sorting Complexes Required for Transport/Endolysosome Pathway and Tau Clearance. Biol Psychiatry. 92:10–24.

Liu, H., A.Y.S. Tan, N.F. Mehrabi, C.P. Turner, M.A. Curtis, R.L.M. Faull, M. Dragunow, M.K. Singh-Bains, and A.M. Smith. 2025. Astrocytic proteins involved in regulation of the extracellular environment are increased in the Alzheimer’s disease middle temporal gyrus. Neurobiology of disease. 204:106749.

Liu, Q., J. Bautista-Gomez, D.A. Higgins, J. Yu, and Y. Xiong. 2021. Dysregulation of the AP2M1 phosphorylation cycle by LRRK2 impairs endocytosis and leads to dopaminergic neurodegeneration. Science signaling. 14.

Macia, E., M. Ehrlich, R. Massol, E. Boucrot, C. Brunner, and T. Kirchhausen. 2006. Dynasore, a cell-permeable inhibitor of dynamin. Dev Cell. 10:839–850.

Martini-Stoica, H., A.L. Cole, D.B. Swartzlander, F. Chen, Y.W. Wan, L. Bajaj, D.A. Bader, V.M.Y. Lee, J.Q. Trojanowski, Z. Liu, M. Sardiello, and H. Zheng. 2018. TFEB enhances astroglial uptake of extracellular tau species and reduces tau spreading. J Exp Med. 215:2355–2377.

McMahon, H.T., and E. Boucrot. 2011. Molecular mechanism and physiological functions of clathrin-mediated endocytosis. Nature reviews. Molecular cell biology. 12:517–533.

Mettlen, M., P.H. Chen, S. Srinivasan, G. Danuser, and S.L. Schmid. 2018. Regulation of Clathrin-Mediated Endocytosis. Annual review of biochemistry. 87:871–896.

Michel, C.H., S. Kumar, D. Pinotsi, A. Tunnacliffe, P. St George-Hyslop, E. Mandelkow, E.M. Mandelkow, C.F. Kaminski, and G.S. Kaminski Schierle. 2014. Extracellular monomeric tau protein is sufficient to initiate the spread of tau protein pathology. J Biol Chem. 289:956–967.

Mothes, T., B. Portal, E. Konstantinidis, K. Eltom, S. Libard, L. Streubel-Gallasch, M. Ingelsson, J. Rostami, M. Lindskog, and A. Erlandsson. 2023. Astrocytic uptake of neuronal corpses promotes cell-to-cell spreading of tau pathology. Acta neuropathologica communications. 11:97.

Nandez, R., D.M. Balkin, M. Messa, L. Liang, S. Paradise, H. Czapla, M.Y. Hein, J.S. Duncan, M. Mann, and P. De Camilli. 2014. A role of OCRL in clathrin-coated pit dynamics and uncoating revealed by studies of Lowe syndrome cells. Elife. 3:e02975.

Narayan, P., G. Sienski, J.M. Bonner, Y.T. Lin, J. Seo, V. Baru, A. Haque, B. Milo, L.A. Akay, A. Graziosi, Y. Freyzon, D. Landgraf, W.R. Hesse, J. Valastyan, M.I. Barrasa, L.H. Tsai, and S. Lindquist. 2020. PICALM Rescues Endocytic Defects Caused by the Alzheimer’s Disease Risk Factor APOE4. Cell reports. 33:108224.

Perea, J.R., E. Lopez, J.C. Diez-Ballesteros, J. Avila, F. Hernandez, and M. Bolos. 2019. Extracellular Monomeric Tau Is Internalized by Astrocytes. Frontiers in neuroscience. 13:442.

Reid, M.J., P. Beltran-Lobo, L. Johnson, B.G. Perez-Nievas, and W. Noble. 2020. Astrocytes in Tauopathies. Frontiers in neurology. 11:572850.

Richetin, K., P. Steullet, M. Pachoud, R. Perbet, E. Parietti, M. Maheswaran, S. Eddarkaoui, S. Begard, C. Pythoud, M. Rey, R. Caillierez, Q.D. K, S. Halliez, P. Bezzi, L. Buee, G. Leuba, M. Colin, N. Toni, and N. Deglon. 2020. Tau accumulation in astrocytes of the dentate gyrus induces neuronal dysfunction and memory deficits in Alzheimer’s disease. Nature neuroscience. 23:1567–1579.

Sheehan, P.W., C.J. Nadarajah, M.F. Kanan, J.N. Patterson, B. Novotny, J.H. Lawrence, M.W. King, L. Brase, C.E. Inman, C.M. Yuede, J. Lee, T.K. Patel, O. Harari, B.A. Benitez, A.A. Davis, and E.S. Musiek. 2023. An astrocyte BMAL1-BAG3 axis protects against alpha-synuclein and tau pathology. Neuron. 111:2383–2398 e2387.

Sidoryk-Wegrzynowicz, M., Y.N. Gerber, M. Ries, M. Sastre, A.M. Tolkovsky, and M.G. Spillantini. 2017. Astrocytes in mouse models of tauopathies acquire early deficits and lose neurosupportive functions. Acta neuropathologica communications. 5:89.

Sturner, E., and C. Behl. 2017. The Role of the Multifunctional BAG3 Protein in Cellular Protein Quality Control and in Disease. Frontiers in molecular neuroscience. 10:177.

Sweeney, N., T.Y. Kim, C.T. Morrison, L. Li, D. Acosta, J. Liang, N.V. Datla, J.A. Fitzgerald, H. Huang, X. Liu, G.H. Tan, M. Wu, K. Karelina, C.E. Bray, Z.M. Weil, D.W. Scharre, G.E. Serrano, T. Saito, T.C. Saido, T.G. Beach, O.N. Kokiko-Cochran, J.P. Godbout, G.V.W. Johnson, and H. Fu. 2024. Neuronal BAG3 attenuates tau hyperphosphorylation, synaptic dysfunction, and cognitive deficits induced by traumatic brain injury via the regulation of autophagy-lysosome pathway. Acta Neuropathol. 148:52.

Szabo, M.P., S. Mishra, A. Knupp, and J.E. Young. 2022. The role of Alzheimer’s disease risk genes in endolysosomal pathways. Neurobiology of disease. 162:105576.

Tsunemi, T., Y. Ishiguro, A. Yoroisaka, C. Valdez, K. Miyamoto, K. Ishikawa, S. Saiki, W. Akamatsu, N. Hattori, and D. Krainc. 2020. Astrocytes Protect Human Dopaminergic Neurons from alpha-Synuclein Accumulation and Propagation. The Journal of neuroscience : the official journal of the Society for Neuroscience. 40:8618–8628.

Van Acker, T., J. Tavernier, and F. Peelman. 2019. The Small GTPase Arf6: An Overview of Its Mechanisms of Action and of Its Role in Host(-)Pathogen Interactions and Innate Immunity. Int J Mol Sci. 20.

Wang, P., and Y. Ye. 2021. Filamentous recombinant human Tau activates primary astrocytes via an integrin receptor complex. Nat Commun. 12:95.

Wong, C.O. 2020. Endosomal-Lysosomal Processing of Neurodegeneration-Associated Proteins in Astrocytes. Int J Mol Sci. 21.

Zaccai, N.R., Z. Kadlecova, V.K. Dickson, K. Korobchevskaya, J. Kamenicky, O. Kovtun, P.K. Umasankar, A.G. Wrobel, J.G.G. Kaufman, S.R. Gray, K. Qu, P.R. Evans, M. Fritzsche, F. Sroubek, S. Honing, J.A.G. Briggs, B.T. Kelly, D.J. Owen, and L.M. Traub. 2022. FCHO controls AP2’s initiating role in endocytosis through a PtdIns(4,5)P(2)-dependent switch. Science advances. 8:eabn2018.

Zhang, J., Z. Jiang, and A. Shi. 2022. Rab GTPases: The principal players in crafting the regulatory landscape of endosomal trafficking. Comput Struct Biotechnol J. 20:4464–4472.

Zhou, Y., and H. Sakurai. 2022. New trend in ligand-induced EGFR trafficking: A dual-mode clathrin-mediated endocytosis model. J Proteomics. 255:104503.

